# Associations between infant gut microbiota and the living environment

**DOI:** 10.1101/2025.05.06.651416

**Authors:** Brandon Hickman, Anne Salonen, Kaija-Leena Kolho, Willem M. De Vos, Katri Korpela

## Abstract

The human gut microbiota is central to health and development of the host, and early life microbiota is affected by a range of factors that can alter the infant’s development for years to come. The role of the external, natural environment in shaping the gut microbiota is still largely unknown. We examined how the environment surrounding the home, in terms of land use, air quality, biodiversity, and traffic and road characteristics associate with the infant gut microbiota in the first two years of life with data from 893 children from the longitudinal birth cohort HELMi. We show that the environment has a minimal overall association with the microbiota development. Air quality explained the largest degree of variation in microbiota composition, while proximity to agriculture appeared to have a similar effect as poor air quality during the first 6 months, while open grey space and agriculture had similar associations after 6 months. The results suggest that the infant gut microbiota is not strongly dependent on the external natural environment, and that the impact of the environment is mostly mediated by exposure to pollution that may affect the host’s immune system and indirectly the gut microbiota.

## Introduction

The gut microbiota plays an important role in health and development in early life. Colonization of the infant gut begins at birth through exposure to maternal fecal bacteria (Korpela et al., 2020), but a number of factors affect the development of the gut microbiota throughout the first years of life, such as antibiotics(Dubourg et al., 2014; Theriot et al., 2016), probiotics(Korpela et al., 2018), diet(Notarbartolo et al., 2022; Ruan et al., 2023), and birthmode(Hickman et al., 2024; Reyman et al., 2019). The role of environmental exposures on the infant gut microbiota, however, is poorly known.

Experiments designed to test the importance of environmental microbiota exposure have shown an increase in gut and skin microbial diversity when subjects were exposed to soil and plant - based material(Grönroos et al., 2019; Hui et al., 2019; Nurminen et al., 2018). However, it should be noted that due to the use of animal dung, the microbial sources of the tested materials included also animal and human faecal matter in addition to any environmental microbes. A study investigating specifically nature exposure have not shown changes in host-associated microbiota in adults(Gascon et al., 2020). Many studies assessing the role of urbanization in gut microbiota are comparative studies of populations in rural and urban areas, reporting largely on how urbanization in terms of changes to lifestyle and diet is associated with the gut microbiota, and do not differentiate the environment from these covariates(Ayeni et al., 2018), or don’t adjust for lifestyle, diet or medical differences between the study groups (Lu et al., 2021).

Environmental microbiota can originate from various sources, but soil represents the largest reservoir. However, the upper soil is aerobic and favors aerobic bacteria, which is essential for plant life, while the human gut is anaerobic. There is little overlap between soil and gut microbiota at both high and low taxonomic levels(Tasnim et al., 2017). The gut microbiota is largely dominated by Bacteroidetes and Firmicutes phyla, while Proteobacteria and Verrucomicrobia dominated the soil samples. A mouse study has shown changes to gut microbiota when soil is introduced to cages opposed to clean bedding, with a higher proportion of Bacteroidetes relative to Firmicutes in soil cages (Ottman et al., 2019). However, mice would eat soil and are known for coprophagy.

We examined the infant gut microbiota in the first two years of life as part of the HELMi (Health and Early Life Microbiota) cohort (Korpela et al., 2019) in the capital region of Finland and identified associations between gut bacterial compositions, and a wide range of environmental variables after adjustment for potentially confounding variables.

## Materials and methods

### Study population and environmental data

This study has received written informed consent from parents/guardians for use of samples and data from the children used in this study. The study was approved by the ethical committee of The Hospital District of Helsinki and Uusimaa, Finland (263/13/03/03 2015) and performed in accordance with the principles of the Helsinki Declaration.

The HELMi cohort(Korpela et al., 2019) is a longitudinal birth cohort from the greater Helsinki metropolitan area. Infants were enrolled during the recruitment period between 2016 and 2018. Only healthy singleton infants born term on gestational weeks 37-42 were part of the cohort. Faecal samples were taken at 3, 6, 13, 26, 39, 52, 78 and 104 weeks. For this study, we use data from 893 infants, with 5371 faecal samples, who did not move homes during their first two years of life and completed metadata surveys.

All environmental variables are defined in Supplementary table 1. Outdoor air pollution data was obtained using the SILAM modelling system(Prank et al., 2010), with a resolution of 0.2 degrees, at the geographic location of the home. We calculated the 12-month annual averages of the 100 atmospheric constituents after birth. Due to high collinearity in the air quality data, we performed a principal components analysis (PCA) on the SILAM data and utilized the first six PC scores in the models. Nomenclature was based on correlations between the PC scores and 100 atmospheric constituents.

Biodiversity zonation data describing the biodiversity of Finnish forests (Mikkonen et al., 2020) was also utilized to capture the environmental biodiversity. Three biodiversity classes were utilized: NAT1, NAT2, and NAT6. NAT 1 is the lowest descriptive version giving only information on the degree of deadwood, exposing areas with lots of trees, tree species, and rare forest environments; NAT 2 adds penalties for forestry operations, taking human actions into account; NAT6 combines all biodiversity-related information and provides a list of the most valuable forest areas and landscapes.

Land use classification was obtained from the Corine Land Cover 2018 at 20m resolution using level 2 groupings, apart from of “Industrial, commercial and transport units”, which was broken down to the level 3 definitions of “Road and rail networks and associated land”, “Port Areas”, and “Airports”. Traffic volume (buffered), proximity to different road types in meters, and the length of roads in meters (buffered), were obtained using the digiroad 2018 dataset(*Digiroad*, 2018). There are 8 different classifications of road types (freeway, motorway, main road, main street, local road, private road, dirt road, pedestrian/cycle path). We utilized the road types 1 through 4 for total road length within each buffer size, and all 8 categories for distance to nearest road by road type.

### Microbiota analysis

Bacterial DNA was extracted from the faecal samples using a previously described bead beating methods(Salonen et al., 2010) (Ambion MagMAX™ Total Nucleic Acid Isolation Kit (Life Technologies)) and KingFisherTM Flex-automated purification system (ThermoFisher Scientific) as previously described(Jokela et al., 2023). The 16S rRNA gene amplicon sequencing was performed using Illumina MiSeq and HiSeq platforms for V3-V4 (primers 341 F/785 R) at the Functional Genomics Unit and Institute for Molecular Medicine Finland, University of Helsinki, Helsinki, Finland. The sequencing reads were processed using R package mare(Korpela, 2016), which relies on USEARCH(Edgar, 2010) for quality filtering, chimera detection, and taxonomic annotation. Forward reads (V3), truncated to 150 bases, were used(Korpela et al., 2016). Reads occurring <50 times were excluded as potentially erroneous. The taxonomic annotation was performed using USEARCH(Edgar, 2010) by mapping the reads to the Ribosomal Database Project (RDP) taxonomy database version 18(Cole et al., 2014), restricted to known gut-associated taxa. Taxonomic annotations were verified using RDP classifier and in cases of disagreement, the Blast annotation was used. Potential contaminants were filtered by removing reads appearing in negative controls (PCR or extraction blanks) in corresponding numbers from all samples. The sequencing depth cut-off was chosen to be 3000 reads after QC to samples collected at 3 months or before and to 5000 paired reads for the remaining samples based on species richness to sequencing depth evaluations.

### Statistical analyses

The read counts were summarised at the family level and the data were transformed into relative abundances by dividing with the total read count. Multivariate analysis of variance was conducted using the adonis2 function in the R package vegan. For this, the relative abundances were log-transformed and the Pearson correlation distances calculated. Each environmental variable was modelled separately at each time point, adjusting for birth mode and intrapartum antibiotic exposure, diet (breastmilk, formula, mixed, with or without solid foods), number of siblings, pets, and type of housing (single-family house, row house, apartment).

The taxon-specific associations were modelled using negative binomial models with the data from the first 6 months combined into one model and the data from the subsequent time points into another model. All environmental variables with a significant association to the overall microbiota composition at any time point were combined with the confounding variables into a single model, which was reduced using AIC to identify a significantly associated set of variables for each taxon. For these models, the environmental data were scaled and centered to be able to compare the estimates of variables measured on different scales.

## Results

The gut microbiota composition was analysed based on 16S rRNA gene amplicon sequencing and summarised at the family level. The overall association between the living environment and the gut microbiota was assessed using multivariate analysis of variance (adonis function in vegan), adjusting for birth mode and intrapartum antibiotic exposure, diet (breastmilk, formula, mixed, with or without solid foods), number of siblings, pets, and type of housing (single-family house, row house, apartment).

After the adjustments, overall poor air quality and traffic-related air quality were the most pervasively associated with the gut microbiota (Fig. 1). The maximum amount of the variance explained by any of the environmental variables was 1.22% (traffic-related air quality at 2 years). In addition, factors related to proximity to roads and bodies of water were often significantly associated with the microbiota composition. Other types of land use variables were less commonly associated. Agricultural space was associated with the microbiota at 9-18 months, and non-agricultural green space at 18-24 months.

**Fig. 1.**
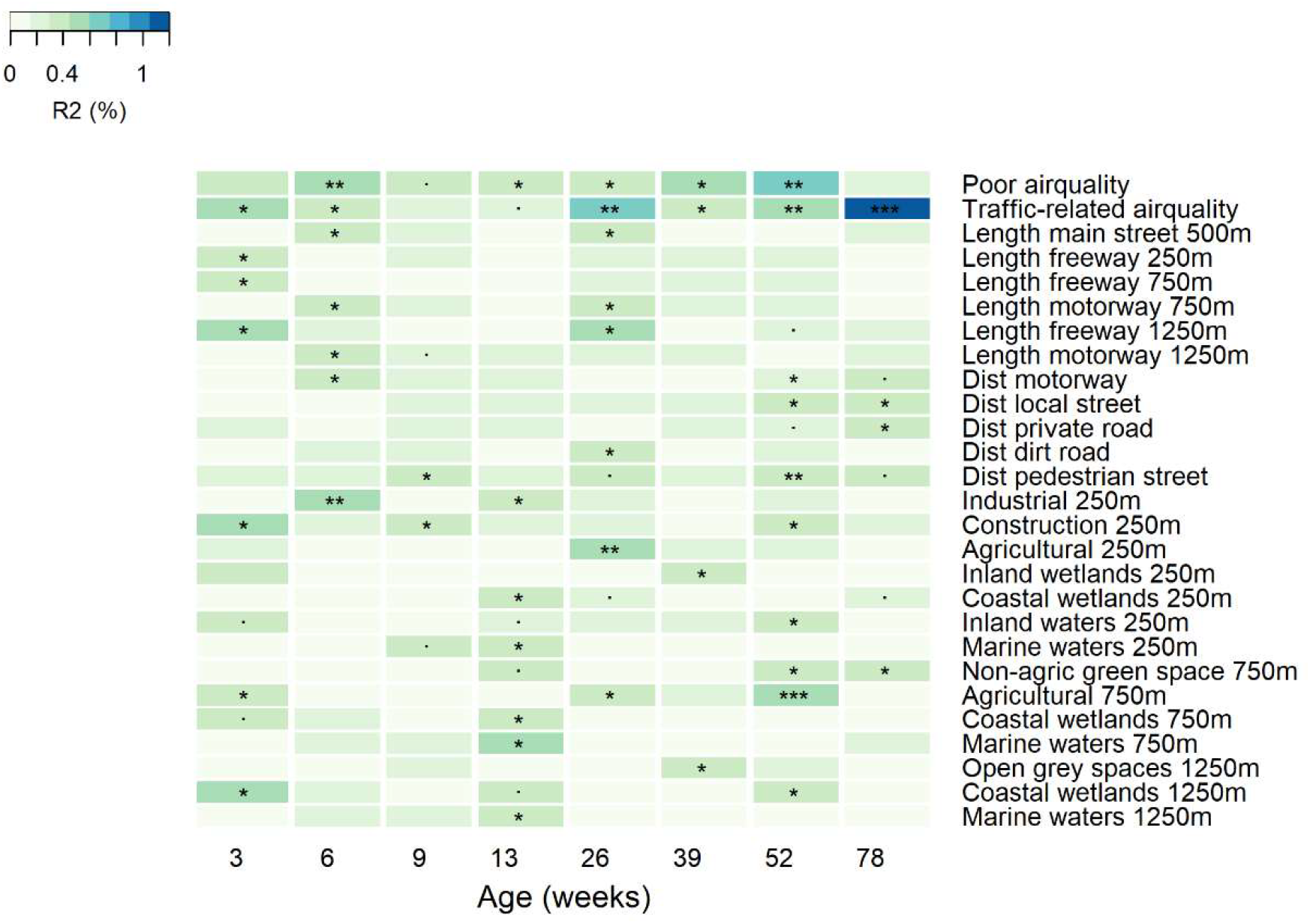
Associations between the gut microbiota and the living environment, based on permutational multivariate ANOVA (adonis). The variance explained (%) is indicated by the color and the asterisks indicate the p-value (., p<0.1; *, p<0.05; **, p<0.01; ***, p<0.001).

To identify the bacterial taxa associated with the environment, the relative abundances of the taxa were modelled against the above-mentioned confounders and the environmental variables which were identified as significantly associated with the overall microbiota composition using generalized linear models. The data were analysed by age, combining samples from the first 6 months into one model and the subsequent time points into another, adjusting for age.

During the first 6 months (Fig. 2), poor overall air quality, distance to a pedestrian/cycle road, motorway, or a private road, agricultural area, inland waters, open grey space, and length of freeway within 1250m buffer (see Supplementary table 1 for variable descriptions) had roughly similar microbiota associations, being mostly positively associated with *Bacteroidaceae, Porphyromonadaceae, Clostridiaceae, Peptostreptococcaceae*, and *Streptococcaceae*, and negatively with *Lactobacillaceae, Erysipelatoclostridiaceae, Micrococcaceae, Eggerthellaceae*, and *Erysipelotrichaceae*. On the other hand, coastal wetlands, industrial/commercial space, and distance to dirt roads, and local streets had similar microbiota associations, being mostly positively associated with *Lachnospiraceae, Eubacteriaceae, Actinomycetaceae*, and *Lactobacillaceae*, and negatively with *Erysipelotrichaceae, Coriobacteriaceae, Porphyromonadaceae*, and *Bacteroidaceae*.

**Fig. 2.**
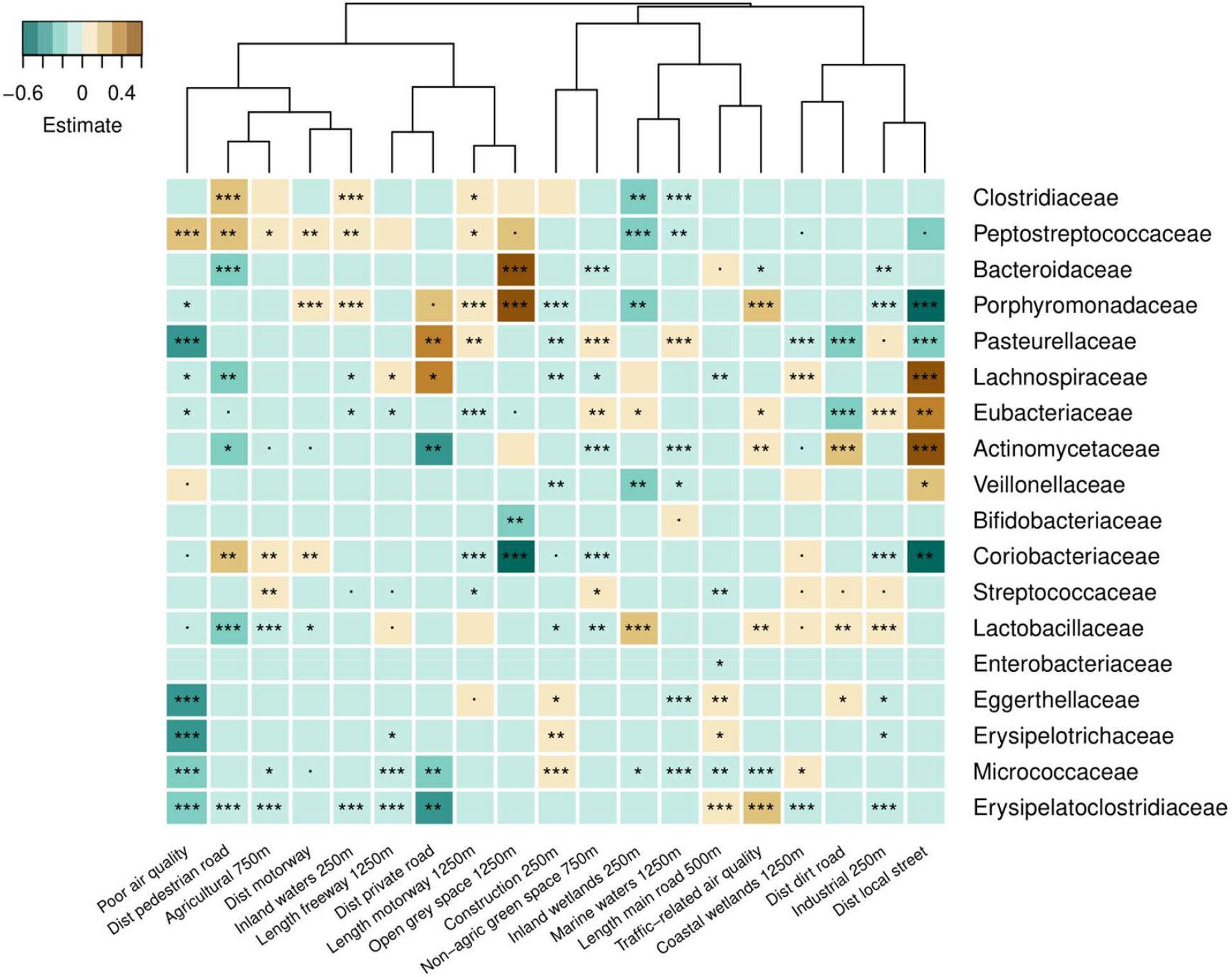
Associations between individual bacterial families in the first 6 months of life and the living environment, based on generalized linear models with the binomial distribution. The model estimate (beta) is indicated by the color and the asterisks indicate the p-value (., p<0.1; *, p<0.05; **, p<0.01; ***, p<0.001).

Between 9 and 24 months (Fig. 3), air quality had the strongest association with the infant gut microbiota composition. Poor general air quality and specifically traffic-related air quality had similar associations as coastal wetlands and distance to nearest private road (private roads are common in rural areas), being positively associated with a large number of taxa, including *Clostridiaceae, Peptostreptoccaceae, Enterococcaeae*, and *Eubacteriaceae*, and negatively with the human milk oligosaccarhides-utilisers *Bacteroidaceae, Akkermansiaceae*, and *Bifidobacteriaceae*. Open grey space and agricultural space had similar associations, mainly a positive association to *Enterobacteriaceae*. Other environmental variables had generally weaker associations.

**Fig. 3.**
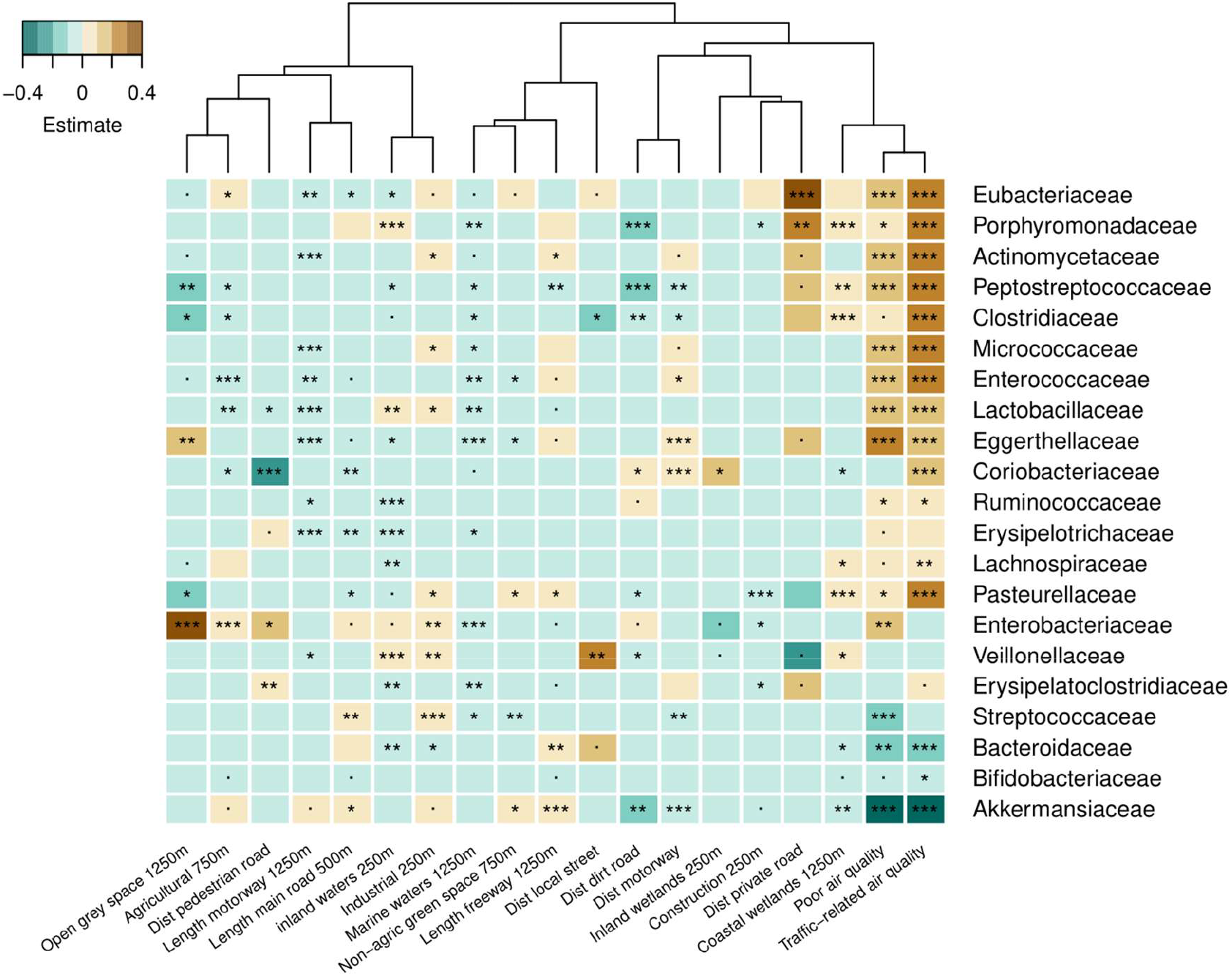
Associations between individual bacterial families at 9 to 24 months and the living environment, based on generalized linear models with the binomial distribution. The model estimate (beta) is indicated by the color and the asterisks indicate the p-value (., p<0.1; *, p<0.05; **, p<0.01; ***, p<0.001).

## Discussion

We found that, after adjusting for important confounders, the surrounding environment has a weak but significant association with the infant gut microbiota composition near the home. Studies that attempt to observe and quantify the impact of the environment on the gut microbiota often lack appropriate confounders, such as diet, ethnic/cultural differences, family history and medical background(Lokmer et al., 2020; Lu et al., 2021; Tyakht et al., 2013). The proximity to forests or forest-related biodiversity near the home, which represent the largest natural environment in Finland covering 67% of the total land area, were not significantly related to the microbiota in any form. The maximum amount of the variance explained by any of the environmental variables was only 1.22%, while we have previously shown with the same data that infant and family-related variables together explain up to 30 % of the variance. Thus, in comparison, the impact of the outside environment on infant gut microbiota is small.

The two air quality variables (poor and traffic-related) were significantly associated to gut microbiota variation throughout the first two years. Exposure to NO_2_, but not residential green spaces, has been associated with a reduction in alpha diversity, as well as changes to the infant gut microbial composition during the first year of life(Cruells et al., 2024). O_3_ exposure has been associated with decreased microbial diversity and a higher abundance of *Bacteroides* spp. in the infant gut (Fouladi et al., 2020). In this study we saw several bacterial families in the infant gut being positively associated with both road related variables and air quality. Significant increases in *Lachnospiraceae, Veillonellaceae*, and *Actinomycetaceae* were seen in conjunction with road related variables such as distance to local streets, and the length of motorways within 1250m of the home and traffic-related air quality. Bailey et al. (2022) saw positive associations between particulate matter with a diameter 10 microns or less (PM_10_,), and the genera *Dialister, Dorea, Acinetobacter*, and *Campylobacter* in the gut microbiota of infants at age of 6 months. Roads and traffic are known contributors to PM that are inhalable into the lungs and can induce adverse health effects. It has been put forward that air pollution may influence the gut microbiota through mechanisms such as gut epithelial damage and permeability, inflammation, and oxidative stress (Van Pee et al., 2023).

Studies on the impact of local environment on the gut microbiota have shown contrasting results. Cross-sectional studies on infants and adults saw that increasing residential greenness levels were associated with a corresponding decrease in gut microbiota diversity(Nielsen et al., 2020; Pearson et al., 2020). However, when comparing residential greenness using vegetation indices from 34 countries and the gut microbiota of adults, an increase in microbial diversity was observed as well as higher relative abundance of *Bifidobacterium* and lower abundance of *Holdemania, Anaerotruncus*, and *Streptococcus*(Zhang et al., 2023). Whether this was in fact mediated by improved local air quality, is not clear, since the study did not adjust for air quality. We found that non-agricultural green spaces, such as urban parks, were negatively associated with *Streptococcaceae*, but only in the first 6 months, after which the association was reversed.

Poor air quality, and agriculture near the home had similar associations to the infant gut microbiota in the first 6 months, while agriculture and open gray space had similar associations after the first 6 months. Thus, proximity to agriculture seems to have the same effect as poor air quality and urban grey space. Modern agriculture practices utilize a variety of pollutants such as pesticide and fertilizers, and there is evidence that people living near fields have an increased exposure than the general population(Dereumeaux et al., 2020). *In-vitro* studies found decreased abundance of *Bifidobacterium* and *Lactobacillus* and increases of *Enterococcus* and *Bacteroides* when exposed to chlorpyrifos (CPF) (Joly et al., 2013; Réquilé et al., 2018; Reygner et al., 2016), the most widely used organophosphate pesticide(Sharma et al., 2023). A similar reduced abundance of *Lactobacilliceae* (first 6 months) and *Bifidobacteriaceae* (6-12 months) was seen in our data.

It was recently reported that infants born during very strict pandemic restrictions, during which infants had very limited exposure to outside environments, had a benign gut microbiota development with a high abundance of bifidobacteria and fewer than expected cases of atopic dermatitis(Korpela et al., 2024), demonstrating that the infant gut microbiota is not strongly dependent on environmental exposure. The degree to which the environment influences the gut microbiota as found in the present study corroborates the findings of Korpela et al. (2024) on the overall independence of the gut from the environmental microbial world. Exposure to the environment seems to influence the gut microbiota mostly indirectly. This is possibly through pollution affecting the host’s immune system rather than directly introducing colonizing microbes. Maternal exposure to air pollution during pregnancy and during lactation has been seen to affect infants after birth and the composition of human milk oligosacrides (Naik et al., 2024). Colonization of environmental microbes would also be difficult as the human stomach has a pH ranging from 1.5 to 3.5, although newborns have a less acidic stomach. Studies have shown that pH <4 results in killing of 99.9% of bacteria within about 90 min(O’May et al., 2005), limiting the potential of microbes on food or from the environment to colonise the gut. It is thus not clear how environmental microbes would plausibly influence the human gut microbiota. Environmental pollution (e.g. PM_2.5_, volatile organic compounds found from indoor gas appliances and exposure to agricultural biproducts, such as pesticides, endotoxins, and allergens) have been shown to upset the immune balance similarly to viral infections(Duramad et al., 2006; Huang et al., 2017; Piao et al., 2023; Zhang & Akdis, 2024). The results suggest that the impact of urbanisation on gut microbiota may be partly driven by increased exposure to pollution, rather than by limited exposure to environmental microbial diversity.(Duramad et al., 2006; Huang et al., 2017; Piao et al., 2023; Zhang & Akdis, 2024)

## Data availability statement

The HELML microbiome 16S rRNA gene sequences in this study have been deposited in the European Nucleotide Archive (ENA) under accession code (https://www.ebi.ac.uk/ena/browser/view/PRJEB55243), along with limited metadata (collection date, sex, age in weeks, geographic location, and sequencing method). Additional individual-level metadata, even pseudonymized, are sensitive and are protected by the GDPR and not publicly available. Reasonable data sharing requests based on data processing and material transfer agreements can be made to Anne Salonen, University of Helsinki, Finland.

## Additional information Funding

This study was supported by the doctoral school of Population Health, University of Helsinki, Finland (B.H.). Microbiota sequencing was partly covered by the SIAM Gravitation Grant 024.002.002, of the Netherlands Organization for Scientific Research (NWO). The study was supported by grants from Tekes 329/31/2015 (W.M.d.V. and A.S.), Academy of Finland 1325103 (A.S.), and Finnish Cultural Foundation (A.S.).

## Notes

https://www.ebi.ac.uk/ena/browser/view/PRJEB55243

### Competing Interest Statement

The authors have declared no competing interest.

